# The Interaction Of Diet-Induced Obesity And Chronic Stress In A Mouse Model Of Menopause

**DOI:** 10.1101/2024.11.11.622997

**Authors:** Nadja Knox, Ali Yasrebi, Daniel Caramico, Kimberly Wiersielis, Benjamin A. Samuels, Troy A. Roepke

**Author notes:** One author has been designated as the corresponding author with contact details: E-mail address, Full postal address: Bartlett Hall, 84 Lipman Drive, New Brunswick, NJ 08901.

## Abstract

Menopause is characterized by the cessation of ovarian hormone production. During postmenopause, cisgender women face increased risks of obesity, cognitive decline, and mood disorder. Mood disorders are associated with exposure to chronic stress. We investigated the combined effects of a high-fat diet (HFD) and chronic stress exposure in a mouse model of menopause using 4-vinylcyclohexene diepoxide (VCD), a selective ovotoxicant that gradually depletes ovarian follicles and hormones. Starting at 6 months, 82 female WT C57BL/6J mice received saline or VCD (130 mg/kg i.p.) 5 days per week for 3 weeks. One month after injection, mice were fed either low-fat diet (LFD) or HFD for 8 weeks followed by 6 weeks of chronic variable mild stress (CVMS). Post-CVMS, mice were either processed for gene expression of the anterodorsal BNST or behavior tests to assess cognitive and anxiety-related behaviors. Plasma samples were collected to analyze metabolic hormones and corticosterone levels. VCD-treated HFD-fed mice had higher fat and body mass, and elevated fasting glucose levels compared to controls and more pronounced avoidance behaviors and cognitive impairments. LFD-fed, VCD-treated mice exhibited less exploration of novel objects and open spaces compared to OIL and HFD counterparts. VCD elevated corticosterone levels on LFD and increased BNST *Pacap* gene expression on HFD. These findings highlight cognitive repercussions of estrogen deficiency and suggest a potential protective effect of a HFD against some of the adverse outcomes associated with menopause. Our study emphasizes the importance of considering dietary and hormonal interactions in the development of therapeutic strategies.

## INTRODUCTION

Menopause is characterized by the cessation of ovarian hormone production. During postmenopause, cisgender women face increased risks of obesity, cognitive decline, and mood disorder (Dalal and Agarwal, 2015; Li et al., 2016). This change has been found to have an increased risk of developing mental health disorders such as anxiety and depression (Bauld and Brown 2009; Freeman et al., 2007). The drop in estradiol has been shown to affect neurotransmitter systems and brain regions involved in mood regulation, leading to heightened susceptibility to mood disorders during and after the menopausal transition. This change has been found to have an increased risk of developing mental health disorders such as anxiety and depression (Bauld and Brown 2009; Freeman et al., 2007; Reis et al., 2014). The drop in estradiol has been shown to affect neurotransmitter systems and brain regions involved in mood regulation, leading to heightened susceptibility to mood disorders during and after the menopausal transition (Reis et al., 2014).

Obesity, prevalent in a substantial portion of the female population, further exacerbates mental health risks, including anxiety and depression. The relationship between obesity and mental health is multifaceted, involving both physiological and psychosocial factors (Patterson et al., 2013; Spencer et al., 2017; Tomiyama, 2018). Excess adipose tissue can lead to chronic inflammation and altered endocrine function, which are implicated in the development of mood disorders (Rosenbaum et al., 2005; Spencer et al., 2017). Obesity and stress are intricately connected, with each exacerbating the effects of the other (Tomiyama, 2018). Additionally, the stigma associated with obesity and the resultant psychological stress contribute significantly to the elevated risk of mental health issues (Puhl and Latner, 2007; Tomiyama, 2018).

Stress disrupts cognitive functions, including self-regulation, which can contribute to unhealthy behaviors such as excessive intake of high-calorie foods, reduced physical activity, and poor sleep quality (Tomiyama, 2018). Chronic stress has been identified as a major risk factor for mental health disorders, including anxiety and depression. The persistent activation of the stress response system leads to dysregulation of the hypothalamic-pituitary-adrenal (HPA) axis, resulting in prolonged exposure to stress hormones like cortisol. This physiological stress response impacts brain function and structure, particularly in regions involved in emotional regulation, such as the hippocampus and amygdala (Kang, 2023). Chronic stress also exacerbates pre-existing mental health conditions and can contribute to the onset of new disorders.

Corticotropin-releasing hormone (CRH) is a key regulator of the hypothalamic-pituitary-adrenal (HPA) axis, the central stress response system. CRH and its receptors are found in various brain regions, including the bed nucleus of the stria terminalis (BNST) (Peng et al., 2017). CRH is released in response to stress and triggers the release of adrenocorticotropic hormone (ACTH) from the pituitary gland, which in turn stimulates the production of stress hormones from the adrenal glands. CRH and other stress-related factors have been implicated in the regulation of both cognition and feeding behavior, with elevated CRH levels being associated with anxiety, depression, and changes in appetite (Hu et al., 2020). Chronic stress and acute activation of the oval bed nucleus of the stria terminalis (ovBNST) lead to maladaptive behaviors in rodents, showing that chronic mild stress activates corticotropin-releasing hormone (CRH) signaling and CRH neurons in the ovBNST in male mice (Hu et al., 2020). Previous studies conducted in our lab have demonstrated that CVMS induces sex-dependent differences in behavioral responses in mice (Hu et al., 2020; Degroat et al., 2022). CVMS exposure resulted in increased anxiety-like behaviors depending on sex and stage of the estrous cycle and have demonstrated that prolonged stress results in cognitive deficits, heightened CRH signaling, increased miniature excitatory post-synaptic activity, and a reduction in M-current activity within the oval nucleus of the BNST in male mice (Hu et al., 2020; Degroat, et al., 2022).

Previous studies have found that corticotropin-releasing hormone (CRH) and associated genes (CRH receptors 1 and 2 (CRHR1 and CRHR2), pituitary adenylate cyclase-activating polypeptide (PACAP), PACAP receptor (PACAPR), and striatal-enriched tyrosine phosphatase (STEP), are involved in the response to chronic stress (Hu et al., 2020; Maita et al., 2022; Lepeak 2023). CRH plays a crucial role in regulating the stress response by stimulating the release of adrenocorticotropic hormone (ACTH) from the pituitary gland. CRHR1 and CRHR2 are the receptors through which CRH exerts its effects, with CRHR1 primarily mediating stress-related responses and CRHR2 involved in modulating the stress recovery process (Baritaki et al., 2019). PACAP and its receptor PACAPR are involved in neuroplasticity and stress responses, influencing both neural and hormonal pathways (Stroth, 2011). STEP has been shown to have a role in regulating the strength of synaptic connections and, consequently, learning and memory processes (Xu et al. 2012; Reinhart et al., 2014).

Therefore, we investigated the combined effects of a high-fat diet (HFD) and chronic stress using a mouse model of menopause - 4-vinylcyclohexene diepoxide (VCD) treatment. VCD, a selective ovotoxicant, gradually depletes ovarian follicles and hormones, mimicking the natural menopausal transition. Unlike ovariectomy, which causes abrupt hormonal loss, VCD provides a more accurate representation of age-related hormonal changes and avoids the stress of surgery. Previous studies have confirmed the ability to induce this level of senescence (Van Kempen et al., 2014; Sui et al., 2023). This experiment builds on our previous VCD studies (Sui et al., 2023), by investigating the combined effects of VCD-induced menopause, HFD, and chronic stress on metabolic outcomes, avoidance and cognitive behaviors, and gene expression in the adBNST. This approach aims to address gaps in understanding how these factors interact and impact physiological and behavioral outcomes, an area not thoroughly explored to date. Our findings seek to increase our understanding of the interactive impact of diet and estrogen levels on physiological and behavioral outcomes and highlight cognitive repercussions of estrogen deficiency and suggest a potential protective effect of a HFD against some of the adverse outcomes associated with menopause. This study provides valuable insights into the mechanisms underlying postmenopausal health risks and emphasizes the importance of considering dietary and hormonal interactions in the development of therapeutic strategies.

## METHODS AND MATERIALS

### Animal Care and 4-Vinylcyclohexene diepoxide (VCD) administration

Procedures adhered to the standards set by the National Institutes of Health and were authorized by the Rutgers Institutional Animal Care and Use Committee. Adult female C57BL/6J mice, purchased from The Jackson Laboratory at 8 weeks of age, were exclusively used in our study. Except for stressors, these mice were housed in a controlled environment with a temperature of 22°C and subjected to a 12-hour light/dark cycle, with unrestricted access to food and water. Mice were administered intraperitoneal injections with either sesame oil (OIL) or 4-vinylcyclohexene diepoxide (VCD) at a dose of 130 mg/kg (in ∼50 mL), administered 5 days per week for a duration of 3 weeks. Body weight was measured every other day (**Figure** 1).

**Figure 1.**
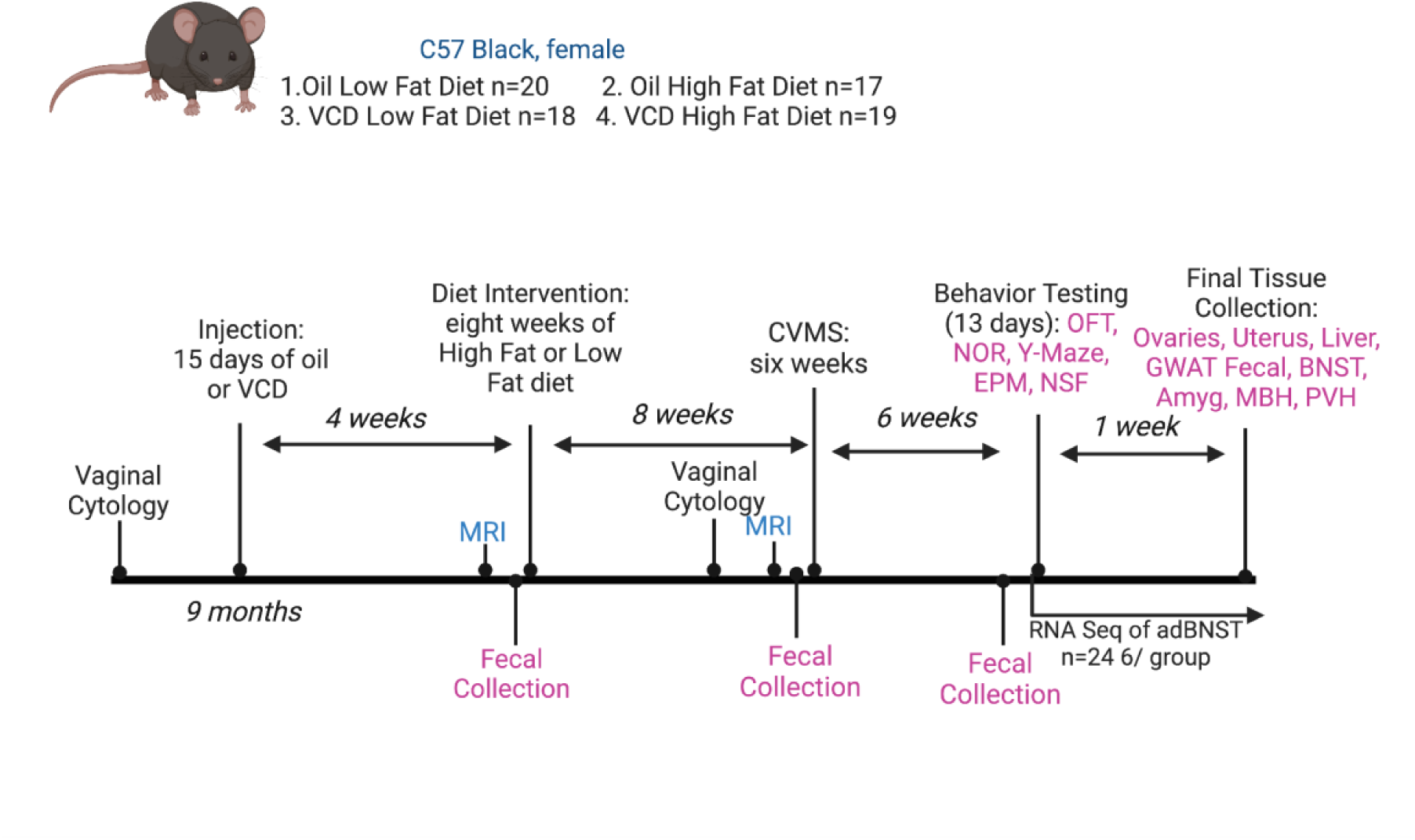
Schematic of experimental timeline. Three cohorts of mice (n=26-28), divided into four treatment groups; OIL LFD (OL), OIL HFD (OH), VCD LFD (VL), and VCD HFD (VH). Administered 130 mg/kg VCD intraperitoneally 5 days per week for 3 weeks, starting at 9 months. Vaginal cytology was performed to classify the stages of the estrous cycle before and after VCD injections. Post-injection, for 8 weeks body weight, food intake, MRI assessments of fat and lean mass content, as well as metabolic parameters were measured. Chronic variable mild stress (CVMS) paradigm was conducted for a six-week period. RNA sequencing as well as qPCR of the anterodorsal bed nucleus of the stria terminalis (adBNST) was conducted post-CVMS to analyze gene expression changes. Behavioral testing (Novel Object Recognition, Y-Maze, Elevated Plus Maze, Novelty Suppressed Feeding), were performed following stress exposure.

### Vaginal Cytology

Vaginal cytology of mice was performed using a lavage technique followed by analysis of the collected cells on a slide, every other day prior to injections, and 14 days straight after injections. This method involved gently flushing the vaginal canal with water, which was then transferred onto a glass slide and spread evenly to create a monolayer. The slide was examined under a light microscope to identify and classify the different stages of the estrous cycle (Caligioni 2009; Cora et al., 2015).

### Diets and metabolic phenotyping

Diet and food intake analysis involved feeding the mice either a Low-Fat Diet (LFD 10% kCal fat, Cat D12450K, Research Diets) or a High-Fat Diet (HFD 45% kCal fat, Cat D12451, Research Diets) *ad libitum* throughout the experiment. Food intake was monitored weekly during eight weeks prior to stress paradigm and behavior testing. The body composition of the mice was assessed using an EcoMRI 3-in-1 Body Composition Analyzer both before and after the dietary intervention. This non-invasive imaging technique collected body composition parameters, such as fat mass and lean mass.

### Chronic Variable Mild Stress (CVMS)

The Chronic Variable Mild Stress (CVMS) paradigm, previously used in published work by our lab, involved subjecting mice to one-two daily stressors over a six-week period. These stressors included alterations in the light/dark cycle (such as overnight illumination), temperature fluctuations (such as 15 min of cold stress), bedding modifications (including removal of bedding, substitution with wet bedding or one centimeter of room-temperature water, or introduction to the bedding of a novel mouse of the same sex), frequent cage changes (5 or 6 times per day), forced swimming (either in 21 °C water for 4 min or 4 °C water for 2 min), isolation stress lasting for 8 hours or overnight, restraint stress for 1 hour, cage tilting at a 45° angle for 8 hours, and exposure to predator sounds for 15 minutes. Stressors were randomized throughout the 6-week period.

### Behavior Testing

Behavior tests took place in the order of Open Field Test, Novel Object Recognition, Y-Maze, Elevated Plus Maze, and finally Novelty Suppressed Feeding. Mice were given 72 hours of habitation to the behavior room prior to testing. Visual cues were placed on the walls and were not moved throughout all the tests. An empty cage was used as a holding cage to keep mice away from visual cues during intertrial delay. The interior area of testing arenas and the holding cage were sanitized by MB10 between each testing trial to minimize olfactory cues.

### Open Field Test and Novel Object Recognition

The NOR test aims to evaluate the ability of mice to recognize and remember novel objects, providing insights into their spatial working and recognition memory abilities. Mice had five consecutive days of habituation during which mice were allowed to explore the empty open field for five minutes. The first day of the NOR is similar to an Open Field Test (OFT). OFT is a behavioral test used to assess anxiety-like behavior and general locomotor activity, where a mouse is placed in a large, open arena, usually square or circular, and allowed to explore freely. On the sixth day, each mouse experienced a five-minute exposure trial where two new identical objects (either two 250-mL glass flasks or two iron brackets) were again presented in two of the four quadrants of the open field, followed by a five-minute intertrial delay. One of the familiar objects was replaced, at random, with a novel object while the second familiar item stayed in the same location as it had been during the exposure trial. The mouse was reintroduced into the open field and recorded exploring for 5 minutes as the test trial. Video footage capturing their exploration and object approach was analyzed using AnyMaze® software.

### Y-Maze

Y-maze test assesses spatial working memory and exploratory behavior by measuring spontaneous alternation behavior in mice navigating through the maze’s three arms. One arm was designated the starting arm, and the other two arms were defined as the known or novel arm for each test which was counterbalanced between mice. For the exposure trial, the mouse was placed at the end of the starting arm, facing toward the center of the maze. After five minutes of undisturbed exploration of the starting arm and known arm, the mouse was put into the holding cage for a five-minute intertrial delay. Then the divider was removed from the novel arm and the mouse was returned to the Y-maze to freely explore all three arms for five minutes. Video footage capturing their exploration was analyzed using AnyMaze® software.

### Elevated Plus Maze

The EPM test assesses avoidance behavior in mice. Mice were introduced into the central area of a plus-shaped maze, positioned facing an open arm, and allowed to explore for a duration of 10 minutes. Video footage capturing their exploration of open and closed arms was analyzed using AnyMaze® software.

### Novelty Suppressed Feeding

The NSF test helps assess avoidance behavior producing a conflict between the desire to eat after fasting and the fear of venturing into a brightly lit novel arena. Mice were subjected to a 24-hour food deprivation period, starting the day before testing, and were subsequently placed in holding cages at 8:00 A.M. on the testing day. Data was analyzed using AnyMaze® software.

### Blood and tissue collection

Animals were euthanized following sedation with ketamine (100 µL of 100 mg/mL) after a one-hour fast. Various tissues were collected including brain, liver, ovaries, and gonadal white adipose tissue (GWAT) were harvested and stored at -80°C for future analysis. Brain slices were then transferred to RNAlater (Life Technologies, Inc., Grand Island, NE, USA) and stored overnight. The BNST was dissected from slices using a dissecting microscope and stored at -80°C for RNA sequencing analysis.

### Peptide hormones

Plasma analysis of metabolic hormones was conducted using the Luminex® MagPix™ multiplex system (Millipore). Plasma samples were prepared by adding, 4-(2-aminoethyl) benzenesulfonyl fluoride hydrochloride (ABESF), to each collection tube. The prepared plasma samples were used for multiplex analysis using Millipore plate #MMHE-44K, for leptin, ghrelin, and insulin.

### RNA Sequencing

RNA samples extracted from the BNST were subjected to RNA-sequencing. Mice were euthanized after a one-hour humid bedding stressor, and brain slices were extracted and stored in RNAlater solution (Invitrogen) and later microdissected to isolate BNST tissue, which was preserved in RNAlater and stored at -80°C until RNA isolation. RNA extraction was performed utilizing the RNAqueous Micro Isolation kit (Invitrogen), and RNA integrity was assessed using an Agilent 2100 Bioanalyzer. All samples had a RNA Integrity Number of 8 or higher. The RNA samples were then sent to the JP Sulzberger Columbia Genome Center (New York, NY) for sequencing. Library preparation was carried out using the TruSeq Stranded mRNA Library Prep Kit (Aviti Element), and the pooled libraries were sequenced on the Illumina NovaSeq 6000 platform with 100 bp paired-end reads, aiming for a sequencing depth of 40 million reads. Subsequently, the resulting reads underwent quality trimming using the FastX Toolkit (http://hannonlab.cshl.edu/fastx_toolkit/) with a minimum quality score of 20.0. Alignment to the mm10 reference genome was performed using STAR along with the NCBI RefSeq mm10 gene annotation, and reads were counted using FeatureCounts.The data were subsequently imported into Rstudio, to conduct differential expression analysis using DESeq2 with the design formula: (diet and Oil/VCD). Principal component analysis (PCA) was performed using Rstudio to identify any outliers.

### Quantitative real-time PCR

cDNA was synthesized from total RNA using Superscript III reverse transcriptase (Life Technologies, Inc), 4 μL 5× buffer, 25 mM MgCl_2_, 10 mM deoxynucleotide triphosphate (Bio-Rad Laboratories, Inc), 100 ng random hexamer primers (Bio-Rad Laboratories, Inc), 40 U/μL Rnasin (Bio-Rad Laboratories, Inc), and 100 mM dithiothreitol in diethylpyrocarbonate-treated water (Gene Mate; Bioexpress, Inc) in a total volume of 20 μL. Reverse transcription was conducted using the following protocol: 5 minutes at 25°C, 60 minutes at 50°C, and 15 minutes at 70°C. The cDNA was diluted to 1:20 with nuclease-free water (Gene Mate; Bioexpress) resulting in a final cDNA concentration of 0.5 ng/μL and stored at −20°C. The adBNST tissue RNA was used for positive and negative controls (no reverse transcriptase) and processed with the experimental samples. Primers were designed and synthesized by Life Technologies using Clone Manager 5 software. For qPCR, 4 μL of cDNA template was amplified using SSO Advanced SYBR Green (Bio-Rad Laboratories, Inc). The amplification protocol followed for all the genes: denaturing at 95°C for 3 minutes followed by 40 cycles of amplification at 94°C for 10 seconds (denaturing), 60°C for 45 seconds (annealing), and completed with a dissociation step for melting point analysis with 60 cycles of 95°C for 10 seconds, 65°C–95°C (in increments of 0.5°C) for 5 seconds and 95°C for 5 seconds. The reference genes used were *GAPDH* and *HPRT*.

### Statistical Analysis

All parameters, unless otherwise specified, were analyzed using Prism (GraphPad Prism) with a significance threshold set at a p-value or adjusted p-value of < 0.05. Prior to analysis, behavioral data was screened for outliers using the Grubbs’ test, employing a critical value of 0.05 as the cutoff. Subsequently, data were subjected to a two-way ANOVA (VCD, diet) followed by Fisher’s LSD *post-hoc* comparison, as we were only interested in comparison within treatment groups (diet or VCD treatments).

## RESULTS

### Metabolic parameters

As metabolic disturbances such as diet-induced obesity may interact with menopause to influence the response to chronic stress (Patterson et al., 2013; Kumari 2023; Shetty et al., 2024), we first characterized metabolic parameters in all groups. HFD-fed, VCD-treated mice displayed a significant increase in body mass gain (Figure 2A), percentage of body weight gain (Figure 2B), and cumulative food intake (Figure 2C) compared to HFD-fed, oil-treated mice. Additionally, fat mass gain (Figure 2D), percentage of fat mass gain (Figure 2E), and body weight difference (Figure 2F) were significantly higher in the HFD-fed, VCD-treated group. Lean mass gain (Figure 2G) and percentage of lean mass gain (Figure 2H) showed no significant differences between the groups. Feed efficiency (Figure 2I) was notably increased in the HFD-fed, VCD-treated mice.

**Figure 2.**
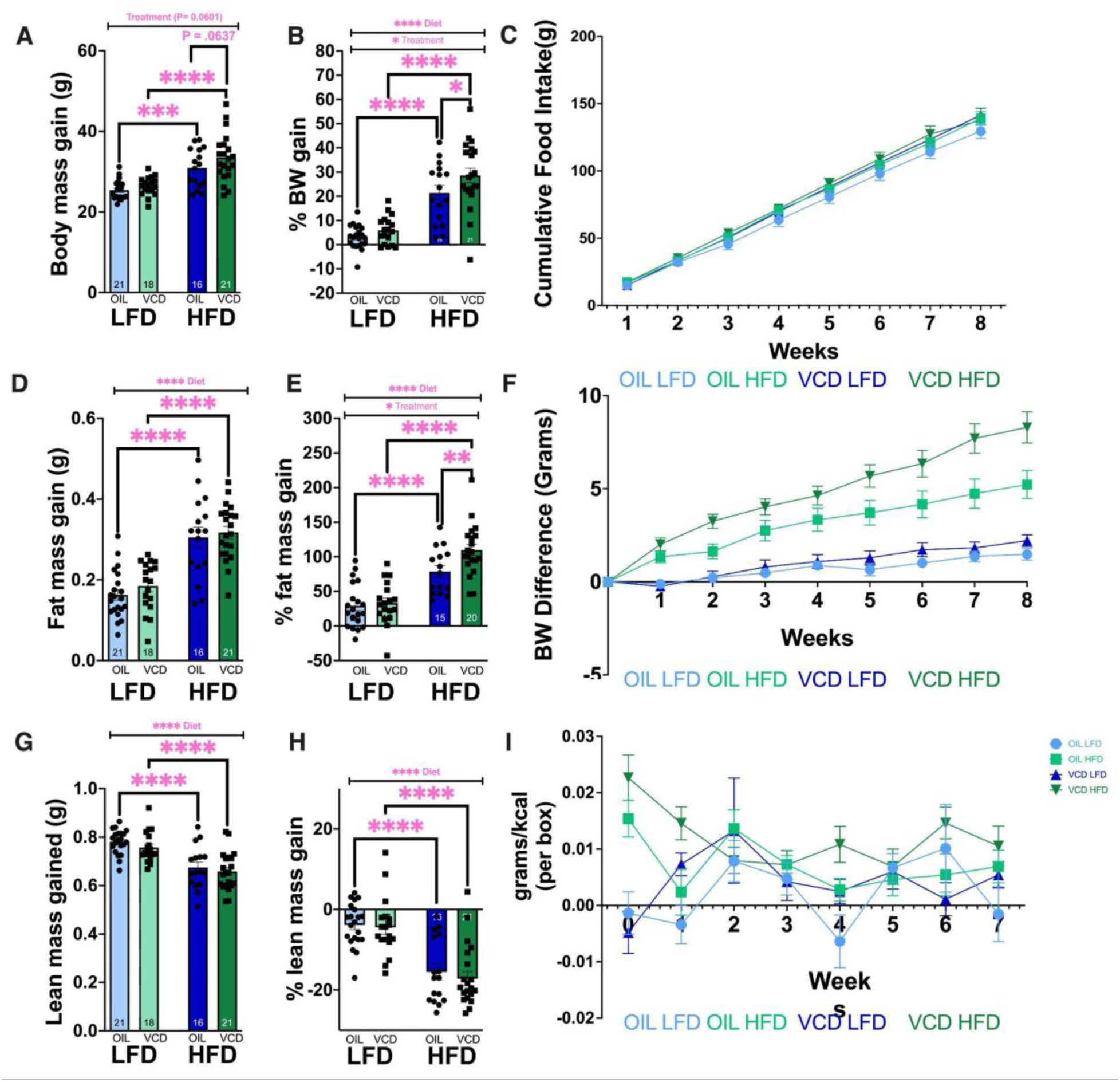
Weight gain, food intake, and body composition in oil- and VCD-treated females fed either LFD or HFD. (A) body mass gain (B) % body weight gain (C) Cumulative food intake (D) fat mass gain (E) % fat mass gain (F) body weight difference (G) lean mass gained (H) % lean mass gain (I) feed efficiency per box. HFD-fed, VCD-treated mice displayed significant increases in body mass gain, percentage of body weight gain, and food intake, compared to HFD-fed, oil-treated mice. Additionally, fat mass gain, percentage of fat mass gain and body weight difference, were significantly higher in the HFD-fed, VCD-treated group. Data presented as mean +/-SEM and analyzed by two-way ANOVA with uncorrected Fisher’s LSD post-hoc comparisons (*, P=0.01 to 0.05, **, P=0.001 to 0.01, ***, P=0.0001 to 0.001).

A two-way ANOVA analysis of cumulative body weight (BW) indicated interaction (F(24,664)=4.204, p<0.0001, *d*=.01), diet (F(8,664)=35.38, p<0.0001, *d*=.36), and VCD treatment (F(3,664)=147.5, p<0.0001, *d*=.03) effects. Weekly body weight fluctuations revealed an effect of VCD treatment (F(3,629)=12.99, p<0.0001, *d*=.05). Cumulative food intake per animal revealed no interaction effect (F(21,275)=0.3485, p=0.9970, *d*=.00). However, there was a VCD treatment effect (F(3,275)=7.767, p<0.0001, *d*=.00). Weekly energy intake per animal calculations showed no interaction effect (F(21,279)=0.6987, p=0.8338, *d*=.04) or time effect (F(7,279)=1.172, p=0.3191, *d*=.02). However, there was a significant effect of VCD treatment (F(3,279)=38.59, p<0.0001, *d*=.28). Examining feeding efficiency per box revealed a interaction effect (F(21,278)=2.103, p=0.0038, *d*=.12), indicating that the effect of VCD treatment on feeding efficiency varied over time. There was no significant main effect of time (F(7,278)=1.618, p=0.1302, *d*=.03). However, a VCD treatment effect was observed (F(3,278)=10.06, p<0.0001, *d*=.08). Post-diet fat content collected via MRI revealed no interaction between diet and VCD treatment effect (F(1,72)=0.09363, p=0.7605, *d*=.00). As expected, a main effect of diet was found, with the HFD group displaying a higher fat percentage compared to the LFD (F(1,72)=65.57, p<0.0001, *d*=.47), confirming that the HFD increased fat accumulation. In contrast, no main effect of VCD treatment was found on post-diet fat percentage (F(1,72)=0.9917, p=0.3227, *d*=.00). The interaction between diet and VCD treatment did not have an impact on body weight at MRI after 8 weeks on specified diet (F(1,72)=0.6333, p=0.4288, *d*=.00). While the difference between OIL and VCD was not significant (F(1,72)=3.649, p=0.0601, *d*=.03), there is a trend of VCD-treated mice weighing more. The unpaired t-test revealed a difference in five hour fasting glucose levels between the VCD and OIL groups (t=2.624, df=25, p=0.0146), with the VCD group having a higher mean value (143.5) compared to the OIL group (124.1).

### Behavioral Analysis

In the OFT, there was a diet effect on distance traveled (F(1,48)=8.271, p=0.0060, *d*=.13) and on distance itself (F(1,48)=8.759, p=0.0048, *d*=.12, Figure 3). The VCD treatment effect was significant for all measured parameters (p<0.005), with HFD-fed, VCD-treated mice exhibiting higher levels of anxiety-like behavior compared to control mice. LFD VCD animals displayed a trend towards increased anxiety-like behavior, although this trend was not statistically significant. In the NOR test, there was a significant VCD treatment effect on distance traveled (F(1,47)=15.63, p=0.0003, *d*=.25, Figure 4), mean speed (F(1,46)=15.86, p=0.0002, *d*=.25), novel object entries (F(1,47)=9.538, p=0.0034, *d*=.15), and time oriented towards center (F(1,46)=9.464, p=0.0035, *d*=.01). Interaction effects were significant for distance (F(1,46)=16.02, p=0.0002, *d*=.01), distance traveled (F(1,47)=15.63, p=0.0003, *d*=.01), and mean speed (F(1,46)=15.86, p=0.0002, *d*=.01). Diet effects were significant for novel object entries (F(1,47)=9.538, p=0.0034, *d*=.09), novel object head entries (F(1,47)=7.587, p=0.0083, *d*=.03), and time oriented towards center (F(1,46)=9.464, p=0.0035, *d*=.01). Mice in the LFD VCD group showed impairment in hippocampal-independent memory compared to control mice, with HFD VCD mice exhibiting a trending impairment in hippocampal-independent memory as well.

**Figure 3.**
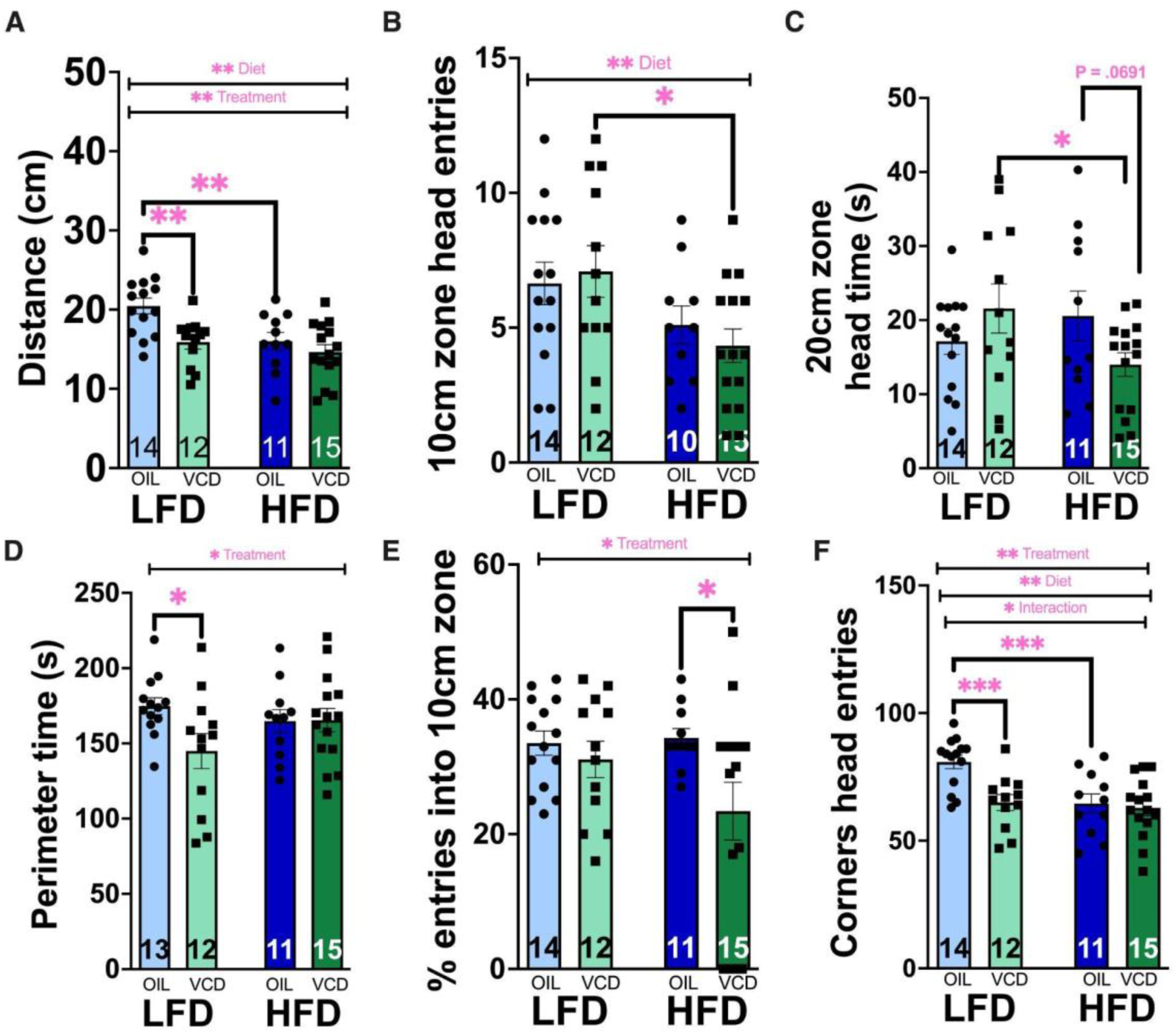
Locomotor activity and avoidance behavior from the open field test in oil- and VCD-treated females fed either LFD or HFD. VCD treatment had a significant effect across (A) distance (B) 10cm zone entries (C) time with head in 20cm zone (D) time in perimeter (E) % entries into 10cm zone (F) head entries into corners. HFD-fed, VCD-treated mice exhibiting increased anxiety-like behavior compared to LFD-fed, oil-treated. Data presented as mean +/-SEM and analyzed by two-way ANOVA with uncorrected Fisher’s LSD post-hoc comparisons (*, P=0.01 to 0.05, **, P=0.001 to 0.01, ***, P=0.0001 to 0.001).

**Figure 4.**
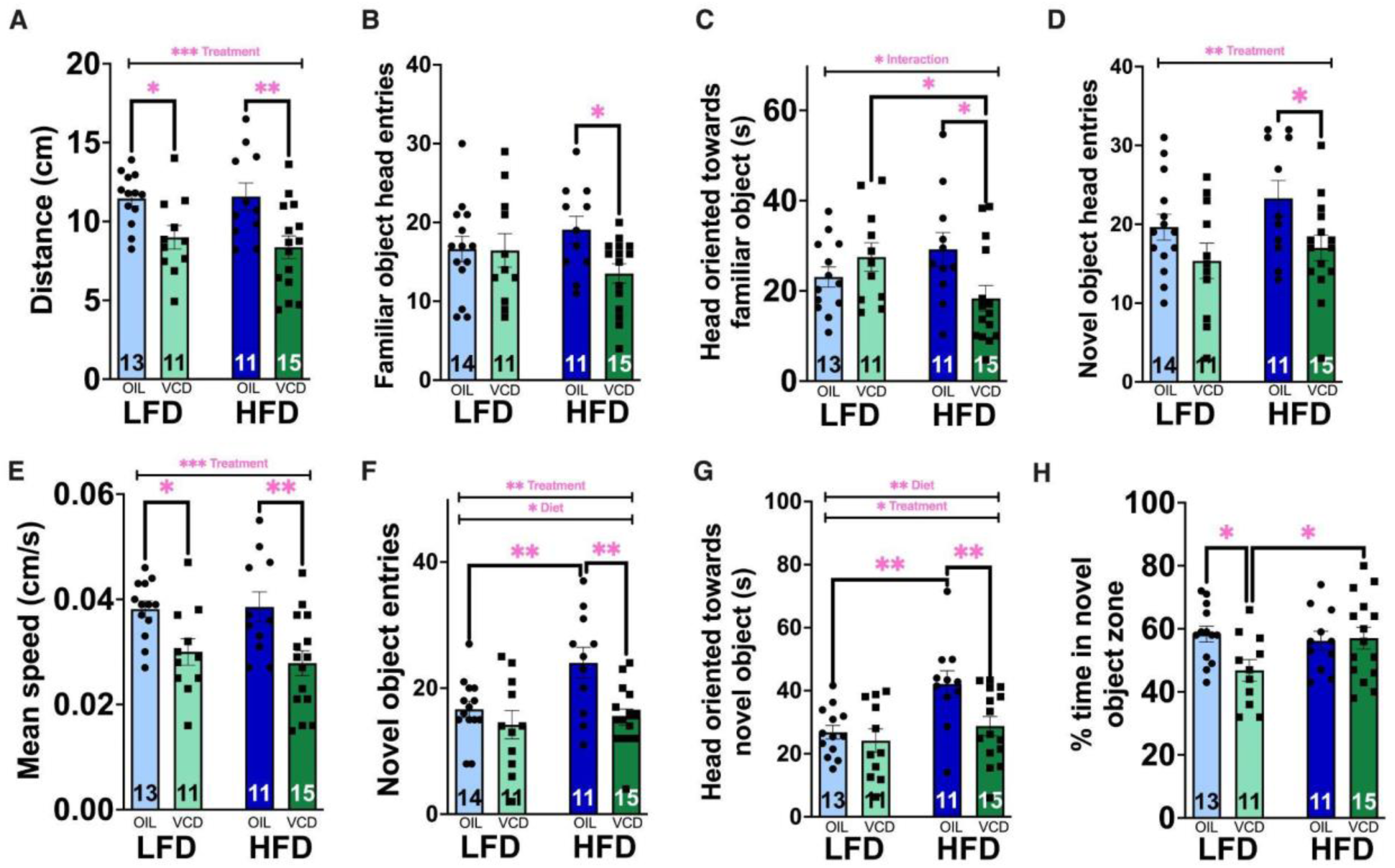
Cognitive and memory in oil- and VCD-treated females fed either LFD or HFD. (A) distance (B) head entries into familiar object since (C) time with head oriented towards familiar object (D) head entries into novel object zone (E) mean speed (F) novel object zone entries (G)time with head oriented towards novel object and (H) % time in novel object zone. VCD treatment impaired memory, with significant effects on distance traveled, speed, and novel object interactions., Data presented as mean +/-SEM and analyzed by two-way ANOVA with uncorrected Fisher’s LSD post-hoc comparisons (*, P=0.01 to 0.05, **, P=0.001 to 0.01, ***, P=0.0001 to 0.001).

In the Y-maze test, there was a significant interaction effect for unknown entries (F(1,48)=10.07, p=0.0026, *d*=.00, Figure 5). Main effects of diet were observed for distance (F(1,48)=6.800, p=0.0121, *d*=.03) and mean speed (F(1,48)=6.945, p=0.0113, *d*=.04). VCD treatment effects were observed for distance (F(1,48)=3.503, p=0.0679, *d*=.12) and mean speed (F(1,48)=3.560, p=0.0658, *d*=.12), indicating differences between oil and VCD-treated animals. Mice fed LFD and treated with VCD exhibited impairment in hippocampal-dependent memory compared to control groups, with HFD VCD mice also displaying a trending decline in hippocampal-dependent memory.

**Figure 5.**
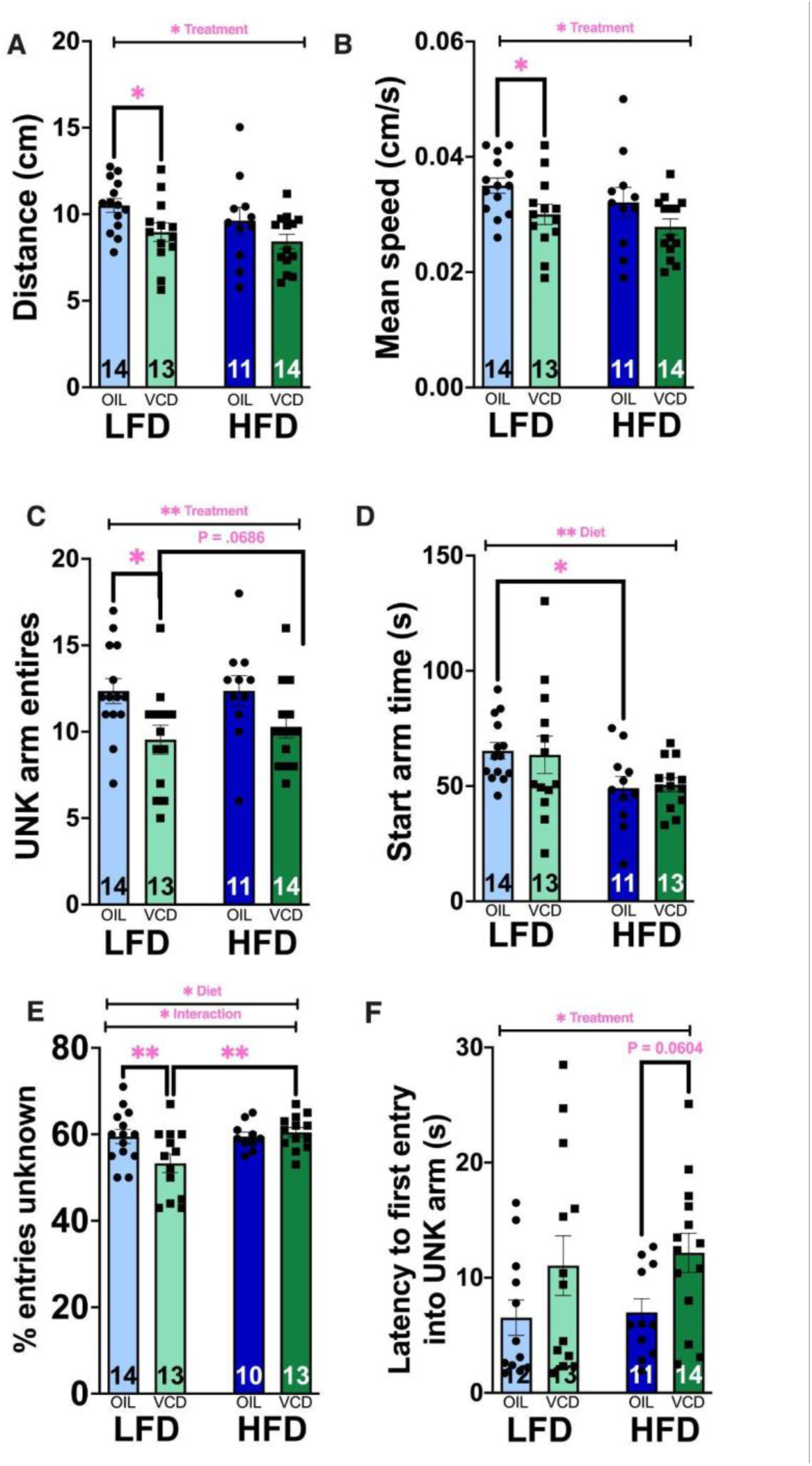
Spatial memory in oil- and VCD-treated females fed either LFD or HFD. (A) distance (B) mean speed (C) unknown arm entries (D) time spent in start arm (E) % entries unknown (F) latency to first entry into unknown arm. VCD-treated mice showed impaired hippocampal-dependent memory, with significant diet and interaction effects on distance, speed, and unknown arm entries. Data presented as mean +/-SEM and analyzed by two-way ANOVA with uncorrected Fisher’s LSD post-hoc comparisons (*, P=0.01 to 0.05, **, P=0.001 to 0.01, ***, P=0.0001 to 0.001).

In the EPM, an interaction effect and a diet effect were found for open arm entries (F(1,42)=3.671, p=0.0623, *d*=.06 and F(1,42)=9.334, p=0.0039, *d*=.03, respectively, Figure 6A and B). Distance and mean speed also showed statistically significant differences between VCD and Oil mice (F(1,42)=3.560, p=0.0658, *d*=.07 and F(1, 44)= 3560, p=0.0658, *d*=.07 respectively). Similar to the OFT findings, mice in the HFD VCD group displayed increased anxiety-like behavior, spending less time in the open arms of the maze. The LFD VCD group also showed a trend towards increased anxiety-like behavior, highlighting a potential VCD treatment effect.

**Figure 6.**
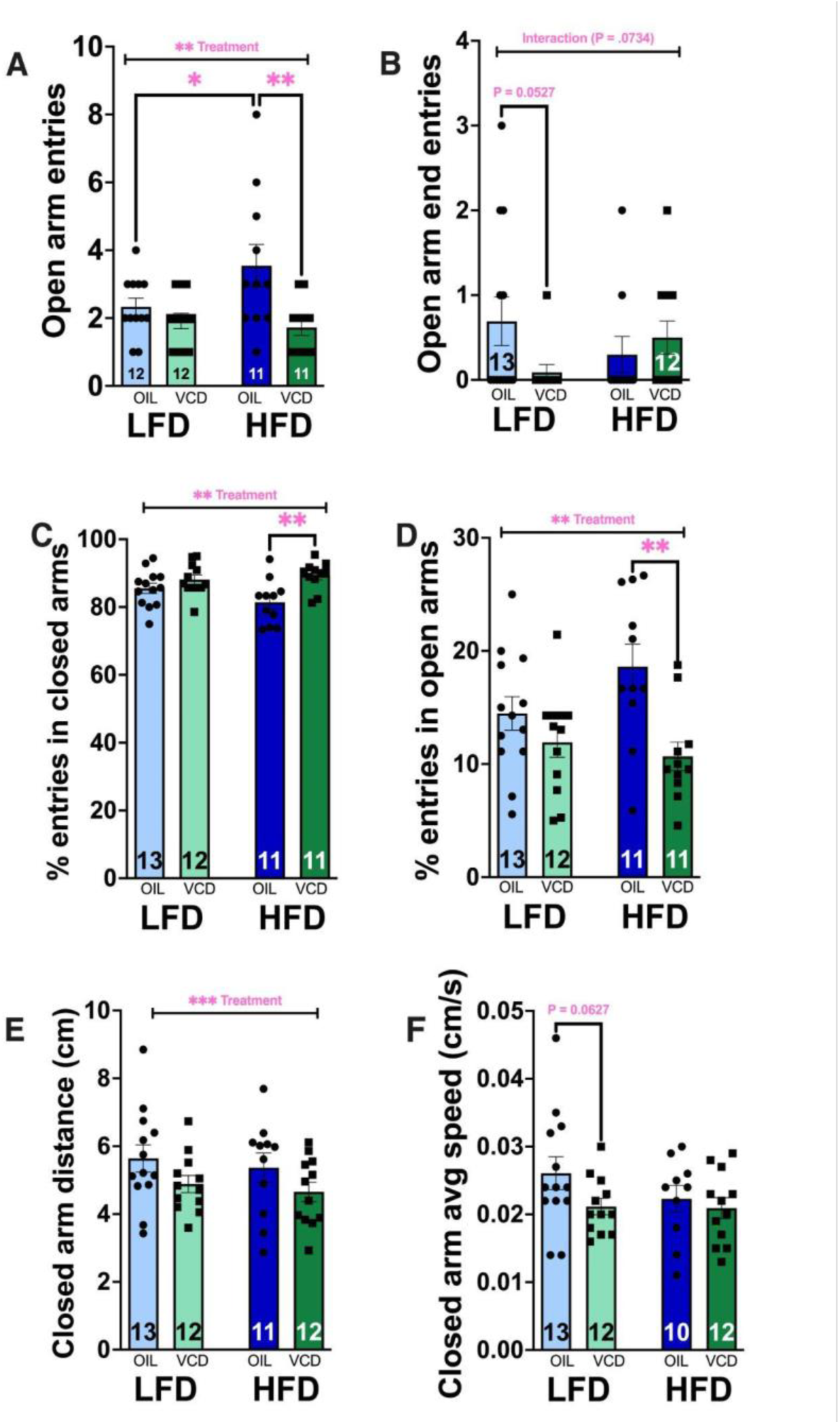
Avoidance behavior in oil- and VCD-treated females fed either LFD or HFD. (A) open arm entries (B) open arm end entries (C) % entries into closed arms (D) %entries into open arms (E) closed arm distance (F) closed arm averages speed. HFD-fed VCD-treated mice showed anxiety-like behavior, spending less time in the open arm. Data presented as mean +/-SEM and analyzed by two-way ANOVA with uncorrected Fisher’s LSD post-hoc comparisons (*, P=0.01 to 0.05, **, P=0.001 to 0.01, ***, P=0.0001 to 0.001).

During NSF testing, only three animals ate within the novel arena, resulting in most animals being disqualified from the study. There were no differences found in latency to eat within the home cage after novel arena testing.

### Peptide hormones

Leptin levels (Figure 7A) were elevated in HFD-fed mice, while VCD-treated mice had higher fasting glucose levels (Figure 7E) compared to oil-treated controls. Insulin (Figure 7D) and CORT (Figure. 7B, 7C) production were influenced by VCD but only in LFD-fed mice. VCD treatment did not have an effect on leptin levels (F(1,34)=2.190, p=0.1482, *d*=.05), nor did the interaction between diet and VCD treatment (F(1,34)=0.0503, p=0.82399, *d*=.00). However, diet had an impact on leptin levels (F(1,34)=7.630, p=0.0092, *d*=.17), with HFD mice having higher levels. Differences were observed between LFD and HFD in the OIL group (p=0.0468). There were no difference in insulin levels based on diet (F(1,32)=0.0045, p=0.9468, *d*=.00), nor VCD treatment (F(1,32)=1.173, p=0.2868, *d*=.03), a trending effect of the interaction between diet and VCD treatment (F(1,32)=3.386, p=0.0750, *d*=.09). Ghrelin levels were not impacted by the interaction between diet and VCD treatment (F(1,33)=0.8787, p=0.3554, *d*=.02) or VCD treatment (F(1,33)=0.0566, p=0.8134, *d*=.00), but there was a trending diet effect (F(1,33)=3.293, p=0.0787, *d*=.09).

**Figure 7.**
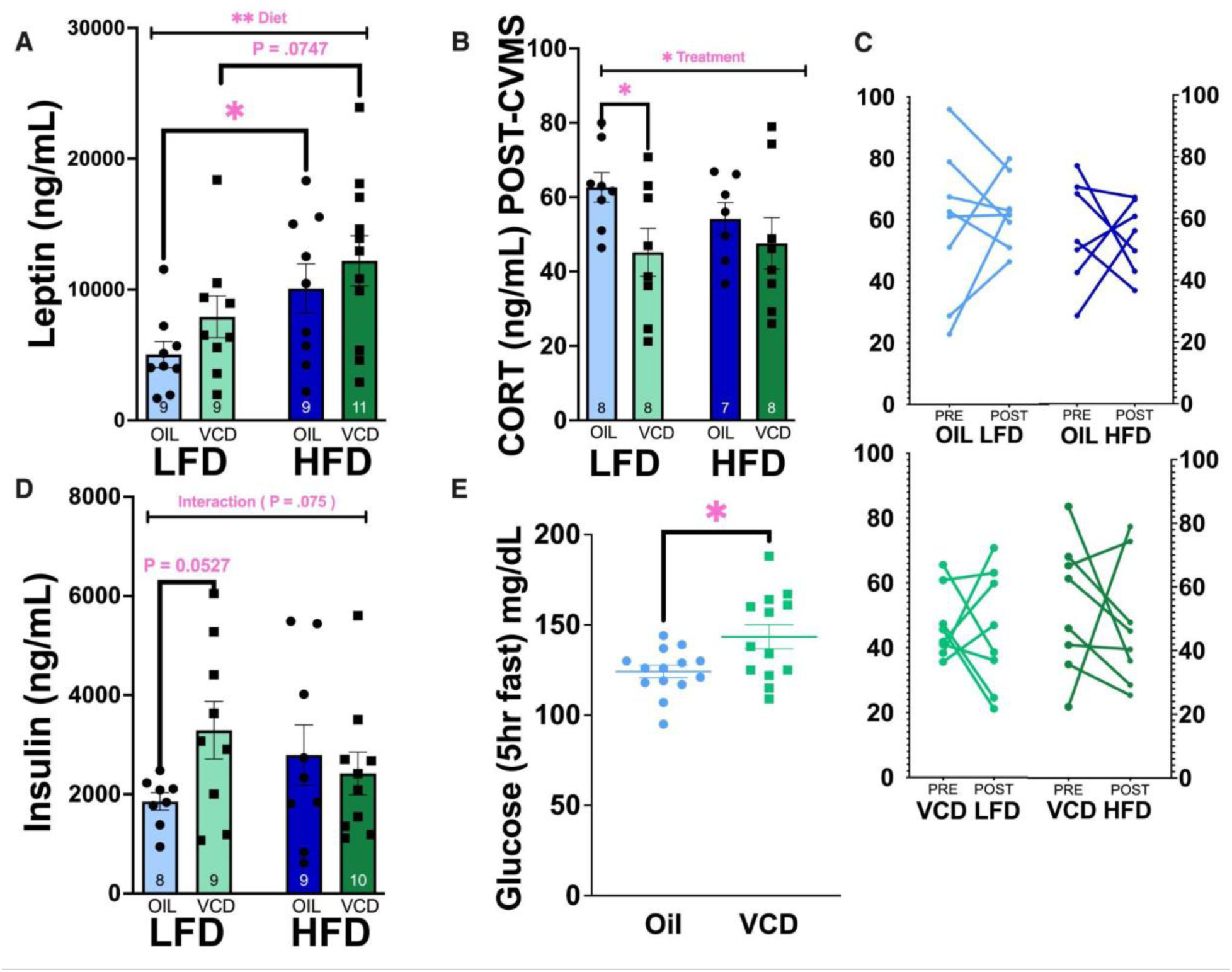
Metabolic and stress hormones in oil- and VCD-treated females fed either LFD or HFD. (A) leptin, (B) CORT, (C) CORT pre and post, (D) insulin and (E) fasting glucose. VCD-treated mice had higher fasting glucose levels, while HFD-fed mice expressed more leptin. Glucose level after a five hour fast were higher in VCD-treated than oil-treated mice. Data presented as mean +/-SEM and analyzed by two-way ANOVA uncorrected Fisher’s LSD post-hoc comparisons (*, P=0.01 to 0.05, **, P=0.001 to 0.01,)

Pre CVMS levels revealed no differences, confirming that corticosterone levels were similar across diet and textural conditions prior to the CVMS protocol, confirming that all mice started from a comparable baseline. Post CVMS levels were not impacted by interaction of diet and VCD treatment effect (F(1,27)=0.9282, p=0.3439, *d*=.03), or diet (F(1,27)=0.2864, p=0.5969, *d*=.01). However, the effect of VCD treatment was significant (F(1, 27)=4.499, p=0.0432, *d*=.14), indicating that variations in treatment conditions (oil vs. VCD) had an impact on corticosterone levels. Post-hoc analyses revealed a significant difference between OIL and VCD under the LFD condition (p=0.0350). No differences were observed for other comparisons. No differences were observed in percent change of levels across the different conditions.

### adBNST transcriptome

RNA-Sequencing and qPCR results for the adBNST are shown in Figure 8. The PCA plot (Figure 8A) and volcano plot (Figure 8B) revealed no significant differentially expressed genes (DEGs) in the transcriptome. While this brain region is hormone and diet sensitive, these findings indicate that chronic stress does not influence the transcriptional response in this region, confirming to what our previous studies have shown (Degroat, et al., 2022). However, PACAP, a CRH activator, was increased by HFD in VCD-treated mice (Figure 8D), while *Crhr1* expression did not show significant changes (Figure 8C). *Crhr1* expression levels did not vary based on interaction between diet and VCD treatment (F(1,18)=2.0155, p=0.1728, *d*=.09), diet (F(1,18)=1.93, p=0.1816, *d*=.09), nor VCD treatment (F(1,18)=1.889, p=0.1862, *d*=.08). However, the comparison between OIL and VCD in the HFD group approached significance (p=0.0748), and the comparison between LFD and HFD approached significance (p=0.0624). Similarly, *Pacap* levels showed no interaction between diet and VCD treatment (F(1,19)=2.275, p=0.1479, *d*=.09), diet (F(1,19)=2.599, p=0.1234, *d*=.10), nor VCD treatment (F(1,19)=0.9765, p=0.3355, *d*=.04) effects. However, the comparison between LFD and HFD within VCD animals was significant (p=0.0357). For *Crh* levels, the interaction between diet and VCD treatment was not significant (F(1,18)=0.7510, p=0.4343, *d*=.01), and neither diet (F(1,18)=0.0830, p=0.7765, *d*=.00) nor treatment (F(1,18)=0.2298, p=0.6374, *d*=.01) had significant effects. We also see this in *Crhr2* levels, the interaction between diet and VCD treatment (F(1,17)=0.5375, p=0.4735, *d*=.02), diet (F(1,17)=2.476, p=0.1340, *d*=.11) nor VCD treatment (F(1,17)=1.683, p=0.2119, *d*=.08) had an effect. For STEP levels, the interaction between diet and treatment was not significant (F(1,17)=0.3428, p=0.5659, *d*=.02), and neither diet (F(1,17)=0.002029, p=0.9646, *d*=.00) nor VCD treatment (F(1,17)=0.05072, p=0.8245, *d*=.00) showed significant effects. For PCAPr1 levels the interaction between diet and VCD treatment was not significant (F(1,18)=0.09983, p=0.7557, *d*=.01) neither diet (F(1,18)=1.015, p=0.3270, *d*=.05) nor treatment (F(1,18)=0.3681, p=0.5516, *d*=.02) had an effect. CRHr1 expression levels showed no significant effects from diet (F(1,18) = 1.93, p = 0.1816, *d*=.08) or treatment (F(1,18) = 1.889, p = 0.1862, *d*=.08), but there was a trend towards significance between OIL and VCD in the HFD group (p = 0.0748) and between LFD and HFD (p = 0.0624). PACAP levels did not significantly interact with diet and treatment (F(1,19) = 2.275, p = 0.1479, *d*=.09), but there was a significant difference between LFD and HFD in VCD animals (p = 0.0357). CRH levels and CRHR2 levels showed no significant effects from diet or treatment. Similarly, STEP and PACAPR1 levels did not exhibit significant effects from diet, treatment, or their interaction.

**Figure 8.**
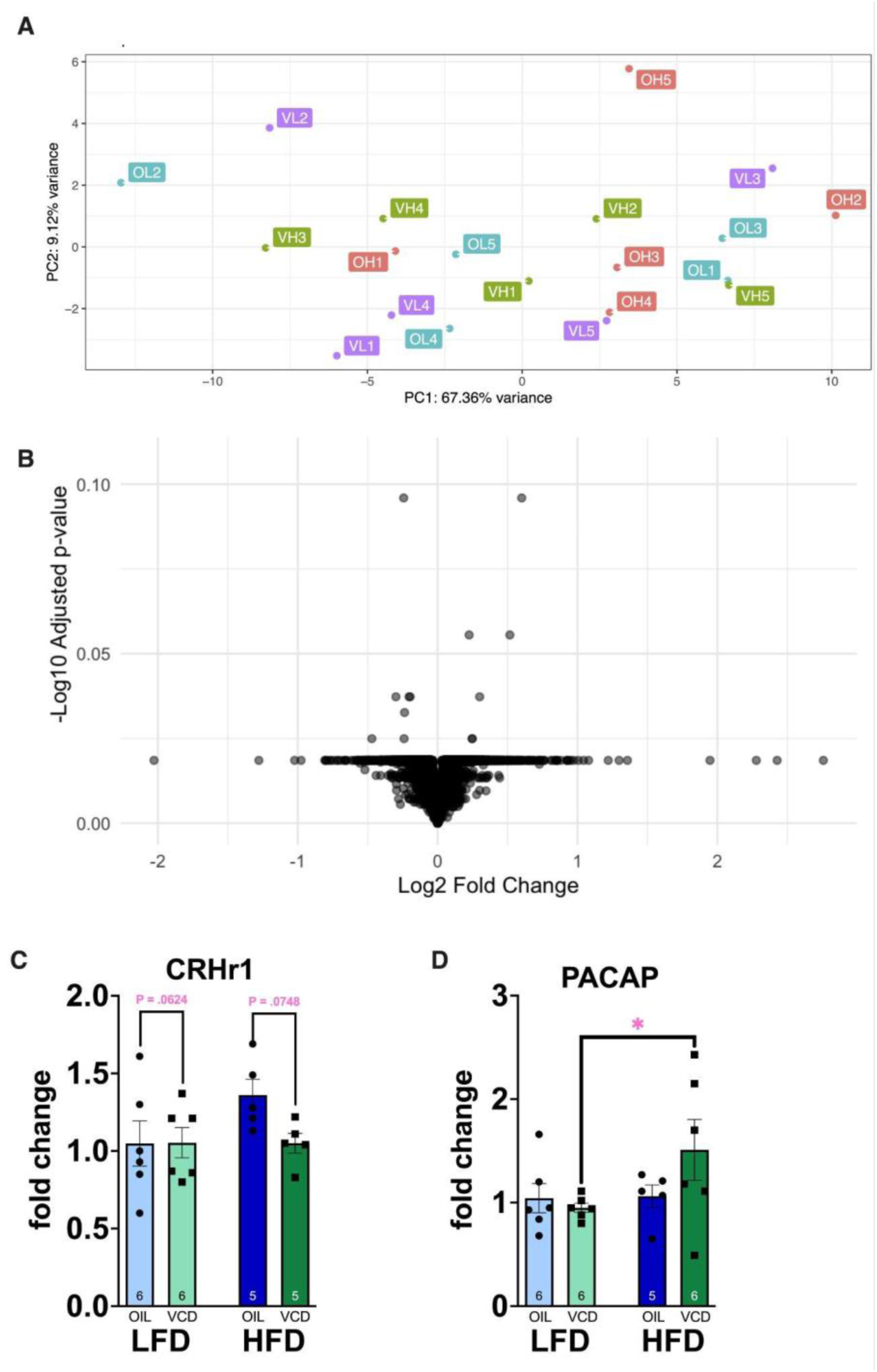
RNA-Sequencing and qPCR of the adBNST in oil- and VCD-treated females fed either LFD or HFD. (A) PCA plot of samples (B) volcano plot (C) *Pacap* gene expression and (D) *Crhr1.* No significant differentially expressed genes (DEGs) were found in the transcriptome of the adBNST, and PACAP expression was elevated in HFD-fed, VCD-treated mice. Data presented as mean +/-SEM and analyzed by two-way ANOVA with uncorrected Fisher’s LSD post-hoc comparisons (*, P=0.01 to 0.05, **)

## DISCUSSION

In this study, we explored the interactions between diet-induced obesity and menopause, in a mouse model of advance ovarian failure by VCD treatment, and their effects on metabolic, behavioral, and transcriptome responses after 6 weeks of chronic stress. We aimed to gain further understanding on how these factors influence anxiety-like behavior, cognition, and hormonal responses, as well as gene expression in the adBNST, a key brain region involved in stress responses. Our results indicate the detrimental impact of HFD and VCD on body composition, anxiety-like behavior, memory impairment, and some metabolic hormone levels in mice. Significant transcriptome changes in the adBNST were not found. We will discuss the metabolic outcomes, followed by behavioral findings, hormonal responses, the transcriptomic data, and future directions.

Body composition changes, including increased body mass gain, fat mass, and feeding efficiency in HFD-fed, VCD-treated mice, align with previous studies linking DIO and menopause to metabolic dysregulation in rodents (Patterson et al., 2013; Kumari 2024). In cisgender women, menopause is associated with increased central adiposity and metabolic syndrome, increasing the likelihood of obesity, largely because hormonal shifts impact energy balance and fat distribution, paralleling our findings in mice (Premaor et al., 2010; Mauvais-Jarvis et al., 2013; Ford et al., 2017). Findings indicate that LFD may increase the risk of weight gain during menopause (Ford et al., 2017). This highlights the importance of dietary suggestions in managing obesity risks and considering potential hormonal changes due to menopause and their impact. In mice, increased central leptin impairs leptin signaling and promotes further weight gain influencing both insulin and leptin regulation (Zhang, 2006). This is also seen when exogenous leptin administration in weight-reduced individuals restores normal energy expenditure and sympathetic nervous system function (Rosenbaum et al., 2005). Leptin and insulin, both elevated in obesity, are known to influence cognitive functions and anxiety (Rosenbaum et al., 2005). In studies studying diet-induced obesity and aging, elevated leptin levels contribute to leptin resistance, which exacerbates obesity (Zhang, 2006). While leptin is primarily associated with energy homeostasis, it also has roles in memory and avoidance behaviors, suggesting that the elevated levels seen in our HFD mice could contribute to the observed cognitive impairments and anxiety-like behavior (Witte, 2016). A previous study demonstrated that VCD treatment induced elevated FSH levels, and impaired hepatic mitochondrial function (Kumari, 2024). Our study further confirms findings that menopause-related metabolic disruptions are exacerbated by a HFD in mice (Sui et al., 2023; Kumari, 2024).

Previous studies on menopause models have reported increased anxiety-like behaviors and impaired avoidance learning, consistent with our findings in the OFT and EPM (Reis et al., 2014; Karisetty et al., 2017; Kang 2023). Administration of estrogen these behaviors, indicating that hormonal changes significantly impact stress-related avoidance and can be mitigated by estrogen (Karisetty et al., 2017; Wada 2018). Our data, showing heightened anxiety in HFD-fed, VCD-treated mice, aligns with these reports and underscores the interaction between metabolic state and menopause. Postmenopausal people are at greater risk for anxiety disorders, particularly when obesity is present, reflecting similar trends seen in our animal model (Bromberger et al., 2013; Karisetty et al., 2017; Tomiyama 2018; Wada 2018). Other research has identified anxiety-like behaviors in the VCD mouse model, showing that treadmill exercise significantly reduces these behaviors and neuronal apoptosis, with the improvements positively correlated with increased levels of circulating osteocalcin (Kang 2023). Our findings differ from previous studies that found HFD impaired fear memory extinction in male mice but not in female mice (Shetty et al., 2024). Our data may differ from other studies due to specific diet composition, duration of diet treatment, duration and type of chronic stress model.

Our results from the Y-Maze and NOR tests indicate that VCD treatment impairs memory, with a more pronounced detrimental impact in HFD-fed mice. These findings are consistent with other studies demonstrating that menopause exacerbates cognitive decline, particularly in memory tasks reliant on hippocampal function (Elsabagh et al., 2007; Bromberger et al., 2013; Premaor et al., 2013; Mauvais-Jarvis et al., 2013; Ford et al., 2017; Wada 2018; Kang et al., 2023). Previous studies have found that while attention, verbal fluency, and memory remain stable, executive function significantly declines in women during late postmenopause, potentially due to hormonal changes (Elsabagh et al., 2007). Intact female mice show better cognitive and emotional resilience compared to ovariectomized mice with more pronounced cognitive deficits and depressive behaviors in response to stress (Karisetty et al., 2017). Alternatively, previous studies have tested conjugated equine estrogens in rats, finding that while it improved cognition in surgically menopausal rats, it did not benefit cognition in transitionally menopausal rats, which more closely mimics the experience of cisgender women (Acosta et al., 2010). Cognitive decline is often reported during the menopausal transition, and obesity is a known risk factor for accelerated cognitive aging (Anacker and Hen, 2017). Adult hippocampal neurogenesis is crucial for reducing memory interference and improving learning and behavioral responses, potentially mitigating anxiety and depressive symptoms associated with aging and menopause (Anacker and Hen, 2017). Again, we acknowledge that variations in our findings compared to other studies could be due to the specific age of at the beginning of treatment, or differences in the timing and type of diet intervention.

High fat diets have been shown to impair cognitive function, avoidance behavior and alter feeding behavior in both humans and animal models (Spencer et al., 2017; Parande et al., 2022; Sui et al., 2023; Kang, 2023). Excessive consumption of high-sugar diets impairs cognitive function and memory, particularly affecting the hippocampus, which is crucial for spatial and episodic memory (Reichelt et al., 2018). Diet induced obesity is widely recognized to negatively impact cognitive function and promote anxiety-like behavior, with mechanisms involving both peripheral and central insulin resistance and altered leptin signaling (Spencer et al., 2017 ; Tomiyama 2018; Parande et al., 2022). Early-life overfeeding and high-fat diets can exacerbate neuroinflammation, impair memory, and increase susceptibility to depressive-like behaviors (Spencer et al., 2017). In the context of menopause, these effects may be magnified, as seen in our VCD-treated, HFD-fed mice. The literature suggests that DIO can exacerbate menopause-related cognitive and emotional disturbances, which aligns with our observations (Van Dijk 2015; Reichelt et al., 2018; Sui et al., 2023). A previous study found that changes in gut microbiota and short-chain fatty acids linked to estrogen levels were associated with cognitive impairments and weight changes, highlighting the connection between menopause, obesity, and cognitive function (Zeibich, 2021).

Our RNA-sequencing analysis of the adBNST revealed no significant DEGs, which agrees with previous studies in stress models where chronic stress does not alter transcriptome in this region (Degroat et al., 2022). This lack of effect may be due to the specific time point we selected for analysis or the possibility that the adBNST’s transcriptional response requires more sensitive detection methods. It may also be due to the fact that in rodents this region is sensitive to stress, all the mice being stressed took precedence over any potential differences due to diet or VCD treatment. A clinical study found that E2 positively influenced cognitive functions such as verbal memory, working memory, and selective attention in healthy postmenopausal women but these benefits were diminished when cortisol levels were elevated due to stress (Baker et al., _2012_). Memory was enhanced by E2 during learning in VCD-treated rats (Koebele et al., 2020). Additionally, E2 alone had a favorable effect on amyloid-β biomarker associated with neurodegenerative diseases, but this effect was counteracted when cortisol was also elevated overall suggesting that elevated stress hormone levels can reduce the cognitive and physiological benefits of estradiol in postmenopausal women (Baker et al., 2012). Previous papers have highlighted that CVMS enhances CRH neuronal excitability in the ovBNST via PKA-dependent CRHR1 signaling (Hu et al., 2020; Maita et al., 2022; Degroat et al., 2023). Blocking CRHR1 or PKA in the ovBNST reversed these electrophysiological and behavioral effects, underscoring the BNST’s pivotal role in the neural circuitry of stress-induced mood disorders (Hu2020). PACAP interacts with CRH signaling in the BNST, with changes in PACAP and CRHR1 expression contributing to stress-induced anxiety-like behaviors (Stroth 2011; Hu et al., 2020; LePeak 2023). However, our data shows an increase in PACAP expression in HFD-fed, VCD-treated mice suggesting that diet may also contribute to changes in stress-related pathways. A study in rats found that PACAP infusion into the BNST increases plasma corticosterone levels (Lezak et al., 2014). The effect was specific to the BNST and was not blocked by CRH receptor antagonism, suggesting PACAP’s key role in stress response regulation (Lezak et al., 2014). CRHR1-deficient mice display increased exploratory behavior and reduced anxiety, underscoring the critical role of CRHR1 in stress and anxiety regulation (Timpl, 1998). The absence of significant changes in CRHR1 expression, despite its role in stress response, also indicates that other mechanisms may contribute to these changes.

A limitation of our study is the absence of unstressed control groups, which would have allowed for further distinction between the effects of diet, menopause, and chronic stress. Future studies should consider the inclusion of some female mice at different stages of the menopausal transition to better model the variability seen in clinical studies and provide context for studies on strategies to manage symptoms of menopause. Other studies can be conducted to look at the different types of stress, such as acute physical stress or social instability stress, to gather further information of the impact of stress.

This study investigates the combined effects of a HFD and VCD-induced menopause on body composition, behavior, hormone levels, and gene expression in mice. HFD significantly increased body and fat mass, particularly in VCD-treated mice. HFD also elevated leptin levels, and VCD treatment raised corticosterone under LFD. Behavioral tests showed that HFD and VCD together exacerbated anxiety-like behavior in the OFT and EPM and impaired memory in NOR and Y-Maze testing. PACAP expression was increased in HFD VCD mice. These findings help provide context to the interactions between diet, menopause, and stress, suggesting that while postmenopausal women with obesity may be at heightened risk for anxiety and cognitive decline, especially if they experience high levels of stress, HFD could offer some protective effect against memory decline. Future studies should consider this information to explore therapeutic strategies to mitigate these risks.

## Summary

The results indicate that HFD-fed, VCD-treated mice had higher fat and body mass compared to HFD-fed, oil-treated mice. Behavioral tests revealed more avoidant behavior and reduced movement in cognitive tests among HFD-fed, VCD-treated mice, implicating the cognitive repercussions of estrogen deficiency. Additionally, hormone analysis showed that VCD-treated mice had higher fasting glucose levels, while HFD-fed mice expressed more leptin. These findings suggest that HFD may increase avoidant behaviors (measures of anxiety) in a menopausal state but highlighting the potential resilience that a high-fat diet provides against some estrogen deficiency-related cognitive impairments such as memory.

## ACNKOWLEDGEMENTS

The authors would like to thank Drs. Judith Storch and Sara Campbell for the use of the EchoMRI™ and Millipore Multiplex Luminex instruments, respectively. This research is funded by NIH/NIMH R01MH123544 (T.A.R. and B.S), USDA-NIFA NJ06195 (T.A.R.), and P30ES05022 (T.A.R.).

The authors declare no competing interests.

